# Millions of Boreal Shield Lakes can be used to Probe Archaean Ocean Biogeochemistry

**DOI:** 10.1101/054478

**Authors:** S. L. Schiff, J. M. Tsuji, L. Wu, J. J. Venkiteswaran, L. A. Molot, R. J. Elgood, M. J. Paterson, J. D. Neufeld

## Abstract

Life originated in Archaean oceans, almost 4 billion years ago, in the absence of oxygen and the presence of high dissolved iron concentrations. Early Earth oxidation is marked globally by extensive banded iron formations but the contributing processes and timing remain controversial. Very few aquatic habitats have been discovered that match key physico-chemical parameters of the early Archaean Ocean. All previous whole ecosystem Archaean analogue studies have been confined to rare, low sulfur, and permanently stratified lakes. Here we provide first evidence that millions of Boreal Shield lakes with natural anoxia offer the opportunity to constrain biogeochemical and microbiological aspects of early Archaean life. Specifically, we combined novel isotopic signatures and nucleic acid sequence data to examine processes in the anoxic zone of stratified boreal lakes that are naturally low in sulfur and rich in ferrous iron, hallmark characteristics predicted for the Archaean Ocean. Anoxygenic photosynthesis was prominent in total water column biogeochemistry, marked by distinctive patterns in natural abundance isotopes of carbon, nitrogen, and iron. These processes are robust, returning reproducibly after water column re-oxygenation following lake turnover. Evidence of coupled iron oxidation, iron reduction, and methane oxidation affect current paradigms of both early Earth and modern aquatic ecosystems.

## Introduction

Ancient oceans on Earth were rich in iron, low in sulfur, and free of oxygen. Oxidation of Earth’s early oceans and atmosphere, concurrent with the early evolution of life and onset of photosynthesis, has long fostered intense scientific debate. Bacterial photosynthesis remains important in modern systems but was key to early Earth oxidation. Only a few modern environments have been identified for whole ecosystem study because the physico-chemical conditions similar to those predicted for the Archaean Ocean are thought to be rare (*1*). Most evidence has been gleaned from the sedimentary rock record and laboratory simulations (reviewed in *2*), spurred by controversy surrounding the origin of the globally ubiquitous and extensive banded iron formations (BIFs).

In a pioneering whole ecosystem study in Lake Matano, Indonesia, large populations of photosynthetic green sulfur bacteria (GSB) were found just below the permanent chemocline at 120 metres depth, including a close relative of *Chlorobium ferrooxidans*, a known photoferrotroph (*3*). This bacterium uses light to fix inorganic carbon, with reduced iron (Fe^2+^) as the electron donor, thereby producing oxidized iron:

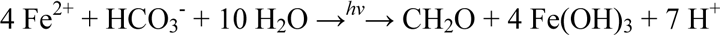

Photoferrotrophy has been proposed as the earliest photosynthetic process in Earth’s history, predating oxygenic photosynthesis by cyanobacteria (*4*). Photoferrotrophy could be responsible for a large part of early Earth oxidation leading to the mixed Fe oxidation states that have proven difficult to explain in globally occurring BIFs, deposited when oxygen was still absent from the atmosphere (*4*,*5*). Since the initial discovery at Lake Matano, only two other ferruginous open water sites (Lac Cruz, Spain (*1*); subbasin of Lake Kivu (*6*), east Africa) have been identified that host microbial communities dominated by photoferrotrophs. Very recently, the first metabolic rate measurements have been reported in the chemoclines of Lakes Kivu and Cruz (*1, 6, 7*), confirming photoferrotrophic activity and its biogeochemical importance by fueling microbial iron reduction and possibly co-existing pelagic heterotrophy. No other examples of such microbial consortia in natural aquatic systems have yet been reported. All of these systems are meromictic and have low sulfate and high iron due to their origins as volcanic craters or rift lakes. Such systems are naturally rare worldwide (*6*), with expectations that only a handful of appropriate lakes will be found (*2*). This limits progress toward understanding Archaean Ocean biogeochemistry and the origins of life.

Almost one half of the largest terrestrial biome on Earth, the boreal forest, is underlain by Precambrian Shield geology (*8*) of low sulfur content. Millions of lakes cover over 7% of the Boreal Shield areas of Canada, Fennoscandinavia, and Russia (*9*). Bottom portions of stratified lakes and shallow ponds become anoxic periodically in summer and under ice in winter if the supply of terrestrial or aquatic organic matter is sufficient and/or the lake morphometry restricts mixing. After the onset of anoxia, the naturally low-sulfur bottom waters of such lakes become rich in dissolved ferrous iron, matching key parameters used in the selection of other Archaean Ocean analogues (*1*). Here we test the hypothesis that the the bottom anoxic layers of seasonally stratified lakes on the Boreal Shield could provide modern *in situ* laboratories for advancing the scientific understanding of microbial metabolic pathways in both the Archaean Ocean and in modern lake environments.

## Results

### Geochemical and stable isotopic evidence

Boreal Shield lakes have been studied intensely at the Experimental Lakes Area (ELA) in northwestern Ontario, Canada. One of these small lakes, Lake 227 (L227), is the site of the world’s longest running nutrient addition experiment, with amendments of nitrogen and phosphorus, or phosphorus alone, for over 47 years (*10*). As a result of fertilization, summer phytoplankton biomass is high, resulting in a hypolimnion that is devoid of oxygen. However, the presence of over 50 cm of continuous annually varved sediments in the deepest part of the lake (10 m; *11*) indicates that the bottom of the hypolimnion has been naturally anoxic during lake stratification for over 300 years under natural nutrient loading conditions. Nearby Lake 442 (L442) has not been manipulated experimentally but the lower portion of the hypolimnion develops anoxia. L442 has physico-chemical features typical of natural lakes on the Boreal Shield. The anoxic zones of both L227 and L442 have low sulfate concentrations (5 to 21 μM) and high total dissolved iron (TDFe) concentrations (115 to 162 μM) over the summer sampling season (Fig. 1), comparable to the levels reported in Lake Matano, Lake La Cruz, and Lake Pavin (*1*). Light penetration into the upper anoxic zones of both lakes is low but detectable, in the range of 0.01-0.03 and 0.09-1.64 μE m^−2^ s^−1^ in L227 and L442, respectively (Supplementary Fig. S1).

**Figure 1.**
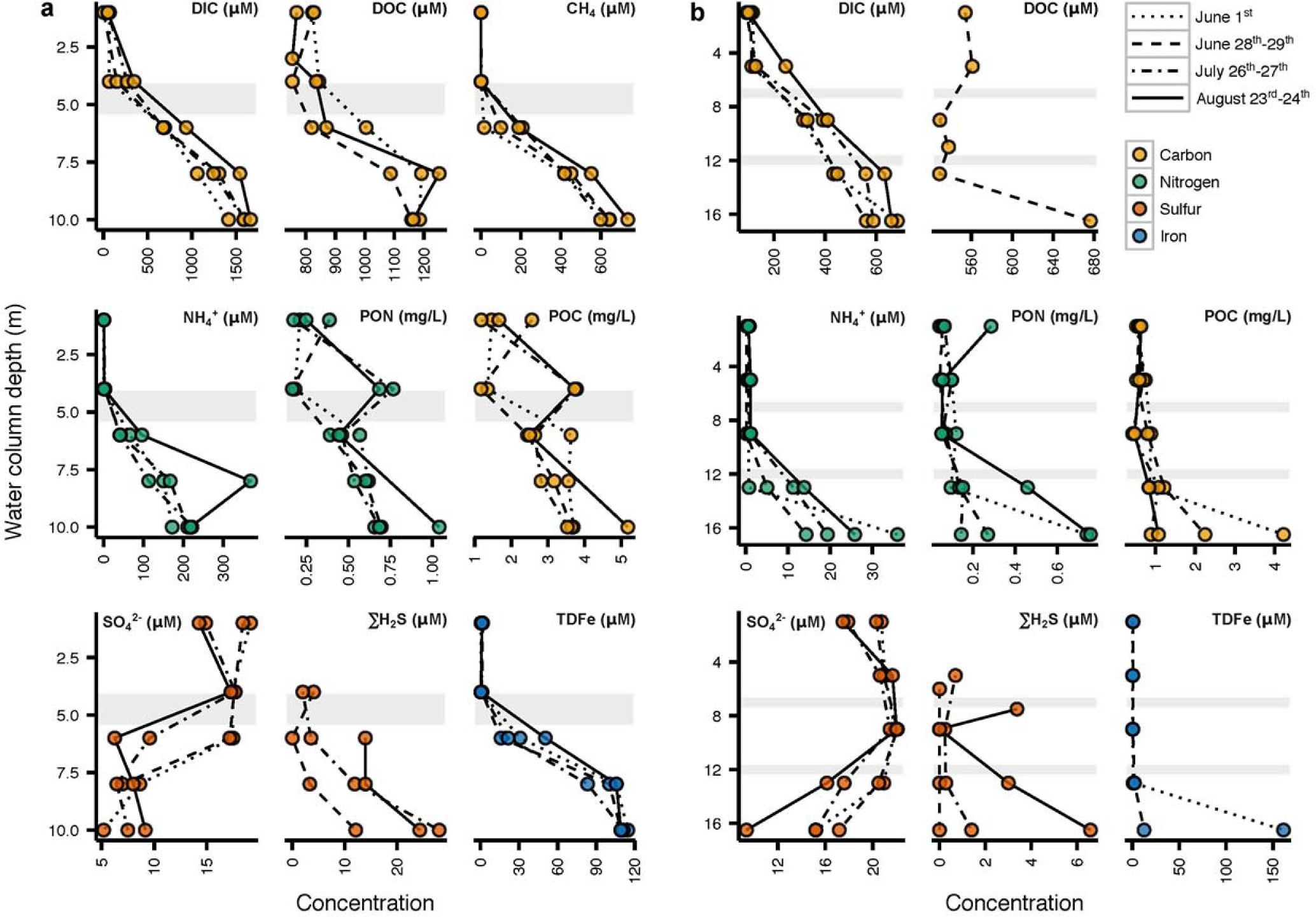
Water chemistry of (a) L227 and (b) L442 water columns. All samples were collected in June-August 2010. Measurements from the same sampling dates are connected by lines for visual clarity. Sampling dates include those from before the onset of the annual cyanobacterial bloom (June 1^st^), during the bloom (June 28^th^-29^th^), and after the bloom (July 26^th^-27^th^, August 23^rd^-24^th^), in L227. In each sub-panel for L227, the surface mixed layer is separated from the seasonally anoxic hypolimnion by a grey transition zone. Each sub-panel for L442 is divided by grey transition zones into the surfaced mixed layer (top), the cool, oxic hypolimnion (middle), and the seasonally anoxic hypolimnion (bottom). The transition zones dividing each lake layer may vary seasonally and annually in both thickness and water column location (depth) due to differing climate (see Supplementary Fig. 2).

Hypolimnetic waters in L227 have distinctive patterns in natural abundance stable isotopes (Fig. 2). These waters are high in iron, ammonium (NH_4_^+^), dissolved inorganic carbon (DIC), and methane (CH_4_), yet low in sulfate (Fig. 1). Stable carbon isotopes (δ^13^C) of particulate organic matter (POM) in the hypolimnion were offset from the overlying epilimnion where phytoplankton biomass was high (Fig. 2a). In contrast, hypolimnetic sediments and sediment trap samples were similar in δ^13^C to epilimnetic POM (Fig. 2a), consistent with the high flux of organic carbon from the surface and indicating that the offset in δ^13^C was not an effect of POM diagenesis during transit through the short water column to underlying lake sediments. Further, δ^13^C of hypolimnetic DIC increased with depth (Fig. 2a). Hypolimnetic POM of lower δ^13^C than either epilimnetic POM or hypolimnetic DIC can only be attributed to isotopic fractionation associated with photosynthesis or assimilation of C with very negative δ^13^C, such as that measured in CH_4_ (Fig. 2a). Similarly, nitrogen isotopic composition (δ^15^N) of hypolimnetic POM was offset from both epilimnetic POM and hypolimnetic NH_4_^+^ (Fig. 2b), consistent with isotopic fractionation during biological uptake of NH_4_^+^ and not N_2_ fixation. Finally, the δ^56^Fe of POM in the epilimnion and hypolimnion (Fig. 2c) also differed and the isotopic fractionation between dissolved and particulate phase was reversed from the epilimnion to hypolimnion. Rates of vertical mixing in the hypolimnion, previously determined in L227 using additions of ^226^Ra and ^3^H as tracers, are very low (*12, 13*), similar to rates of molecular diffusion. These low mixing rates and the persistence over the stratification period mean that microbiota contributing to hypolimnetic POM are suspended in the water column at the observed depth. Together, these data imply that the microbial consortia in the anoxic zone are metabolically active and sufficiently abundant to alter the isotopic POM signatures of δ^13^C, δ^15^N, and δ^56^Fe in POM from values typical of the epilimnion. Furthermore, the offset of the carbon and nitrogen stable isotopes in POM from inorganic dissolved substrates is consistent with fractionation associated with photosynthesis, despite extremely low light levels in the hypolimnion (Supplementary Fig. S1) due to high summer epilimnetic biomass.

**Figure 2.**
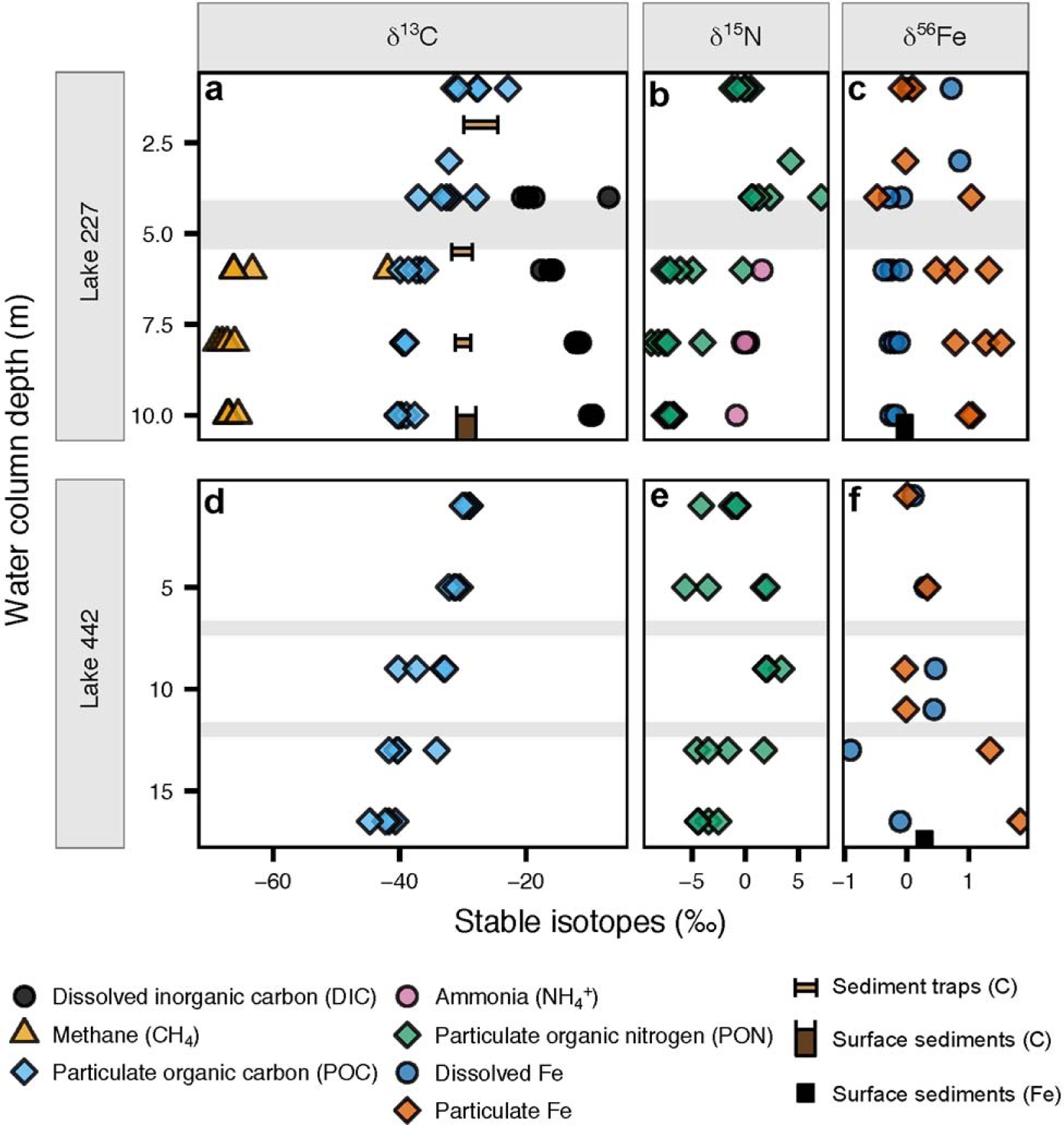
Stable isotopic values in the water columns of (a-c) L227 and (d-f) L442. (a, d) δ^13^ C in DIC, CH_4_, and POC, sampled in June-August 2010 and July 2014. δ^13^ C in sediment traps and surficial sediments (1-2 cm) is also shown. The width of plotted sediment markers represents the range of δ^13^ C values measured across the summer of 2011. (b, e) δ^15^N in NH_4_^+^ and PON, sampled in June-August 2010 and July 2014. (c, f) δ^56^ Fe in dissolved and particulate Fe sampled in June-August 2011 and July 2014. δ^56^ Fe in surficial sediments from July 2010 is also shown. In each L227 panel, the surface mixed layer is separated from the seasonally anoxic hypolimnion by a grey transition zone. Each panel for L442 is divided by grey transition zones into the surfaced mixed layer (top), the cool, oxic hypolimnion (middle), and the seasonally anoxic hypolimnion (bottom). The transition zones dividing each lake layer may vary seasonally and annually in both thickness and water column location (depth) due to differing climate. The water column of L227 at 4 m depth varied substantially in oxygen status over the sampling dates (see Supplementary Fig. 2), which may be responsible for the wide range of isotopic values measured at this depth.

### Microbial community composition

Throughout the L227 hypolimnion, we detected microbial communities that were dominated by iron-cycling bacteria, along with populations of sulfur and methane cycling bacteria, consistent with microbial consortia reported in Lake Kivu, where photoferrotrophic activity was found (*6*). High-throughput sequencing of bacterial 16S ribosomal RNA (16S rRNA) genes from L227 water column samples (Fig. 3) showed *Chlorobi*, closely related to *C. ferrooxidans*, among the most abundant operational taxonomic units (OTUs) in the anoxic zone. Sequences classified within the genus *Chlorobium* comprised up to 8% of all reads just below the oxic-anoxic interface. One particular *Chlorobium* OTU (Fig. 3, *Chlorobium* OTU 1), which was the most abundant across all L227 samples, had over 99.5% 16S rRNA gene sequence identity with known *C. ferrooxidans* strains (Supplementary Fig. S3).

**Figure 3.**
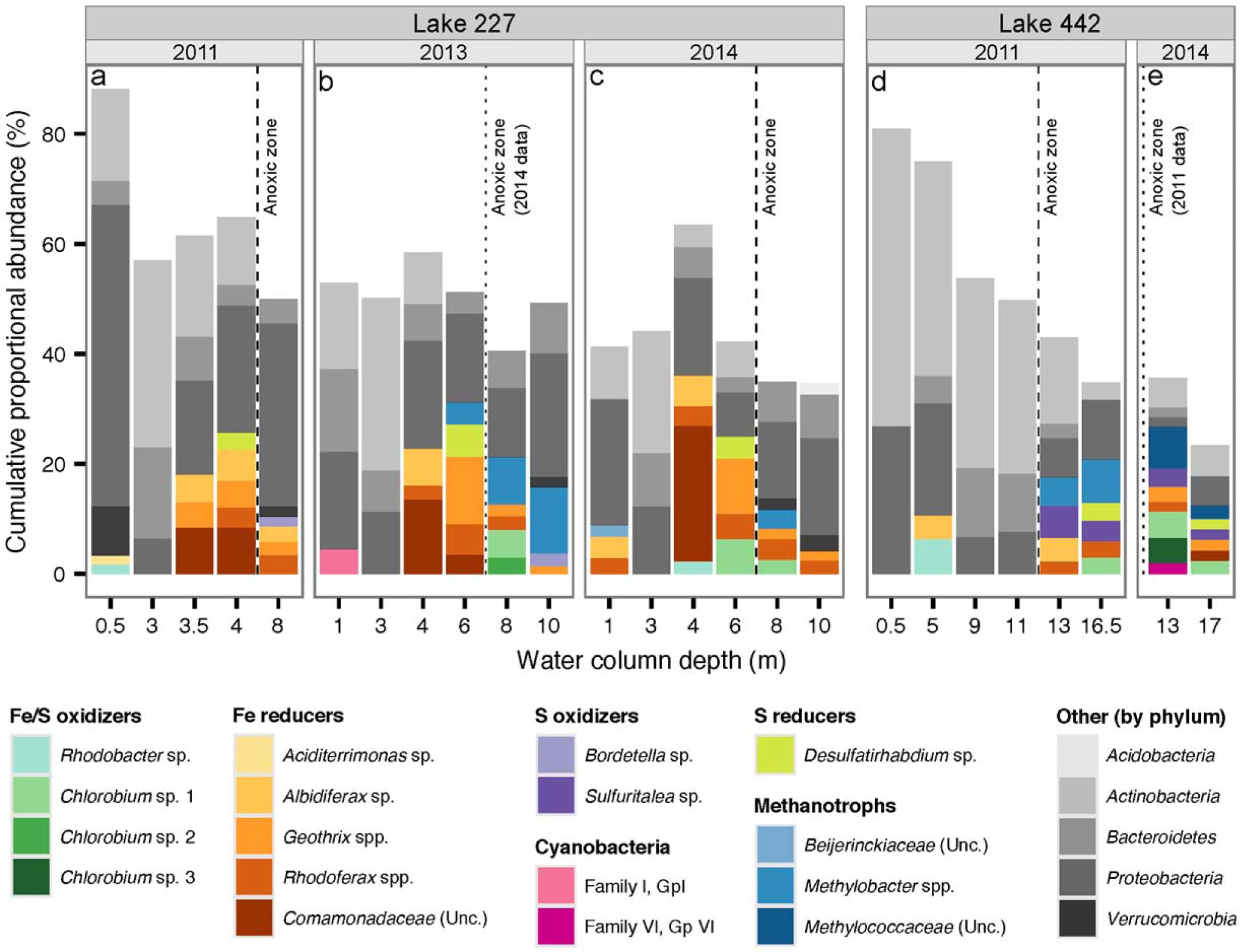
Microbial community in the water columns of (a, b, c) L227 and (d, e) L442. In each depth sample, the ten most abundant bacterial operational taxonomic units (OTUs) are shown as stacked bars in terms of their proportional abundance within rarefied molecular sequencing data. Bacterial OTUs are grouped according to their potential involvement in Fe cycling, S cycling, and methanotrophy (see Methods). Cyanobacterial OTUs are also shown. For clarity, OTUs not associated with these metabolic roles are displayed at the phylum level. Potentially photoferrotrophic *Chlorobium* OTUs are shown with a naming and colour scheme matching that of Supplementary Fig. 3. Although not shown, *Chlorobium* OTU 1 is also present in the anoxic water column in L227 at 6 m depth in 2013 (rank 13, 2.9%) and in L442 at 13 m depth in 2011 (rank 31, 0.67%). All samples were collected between June 25^th^ and July 9^th^ in their respective sampling year.

**Figure 4.**
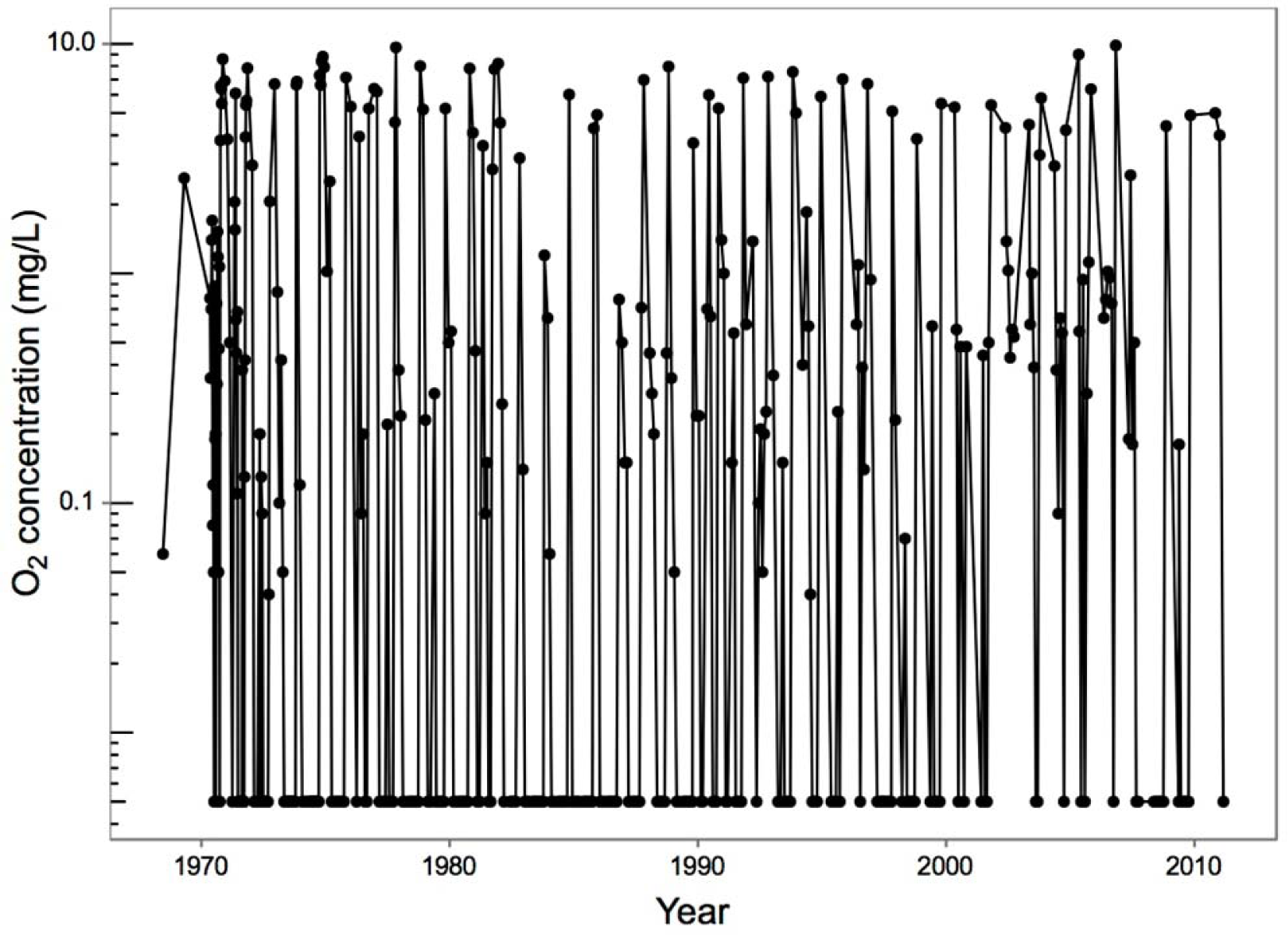
Dissolved oxygen at 6 m depth in the L227 water column from 1969 to 2011. The detection limit for O_2_ is 0.005 mg O_2_ L^−1^. *Chlorobium* sequences were detected at high abundance at 6 m depth in both 2013 and 2014 (Fig. 3, please also see legend). Dissolved oxygen samples were collected typically at least every two weeks in summer but were collected at most twice during the winter. Following this sampling schedule, the full extent of the typical spring and fall re-oxygenation events (overturns) in L227 may not have been measured in some years. Dissolved oxygen is typically, but not always, measured after fall overturn. Spring overturn measurements can be missed following ice-off due to logistical reasons and especially in years when temperatures warm rapidly after ice-off. Thus, the oxygen record at 6 m reflects the minimum number of re-oxygenation events at this depth.

Manually curated functional predictions from 16S rRNA gene data suggest that the hypolimnion of L227 supports a metabolically unique bacterial community. The hypoxic and anoxic zones of L227 were rich in bacterial iron reducers, including members of the genera *Rhodoferax, Geothrix*, and *Aldibiferax*. The OTUs associated with iron reducers were abundant just above and where *Chlorobi* OTUs were identified, implying that these bacteria reduce ferric iron produced by photoferrotrophic activity and chemical oxidation. Putative methanotrophic OTUs belonging to the family *Methylococcaceae* of the *Gammaproteobacteria* were abundant in the anoxic zone. In contrast, potential S-oxidizers and reducers, such as *Sulfuritalea* and *Desulfatirhabdium* spp., were lower in abundance, implying relatively low contributions to overall lake metabolism. Similar populations of *Chlorobium, Rhodoferax*, and *Methylobacter* (family *Methylococcaceae*), along with other sulfur and iron cyclers, were reported in Lake Kivu, the only Archaean Ocean analogues whose whole microbial community has been described to date (*6*). Evidence from targeted sequencing studies in other Archaean Ocean analogues suggests that a similar microbial consortium may be present. In Lake La Cruz, which is known to host photoferrotrophic *Chlorobi*, populations of methanotrophs belonging to the *Alpha*- and *Gammaproteobacteria* were recently identified in the anoxic zone using quantitative polymerase chain reaction (qPCR; *14*). Evidence of anaerobic methane oxidation, perhaps mediated by archaeal methane oxidizers, was found in Lake Matano near the same depth where large *Chlorobi* populations were detected (*15*). Although the presence of photoferrotrophy has not been evaluated in Lake Pavin, sulfur-cycling microorganisms have been identified in the water column, along with populations of bacterial methanotrophs belonging to the genus *Methylobacter*, in the anoxic zone of the lake (*16*). To our knowledge, the distinctive pelagic microbial consortium we identified of iron-cycling bacteria, sulfur-cycling bacteria, and methanotrophs has not been reported outside of Archaean Ocean analogue systems.

### Natural abundance stable isotopes of iron

Key evidence of potential photoferrotrophy is provided by stable iron isotopes, a tool only very recently applied to freshwater lake iron cycling (*17, 18*), and not yet in Lakes Matano or Kivu. In the anoxic hypolimnion of L227, δ^56^Fe in suspended particulates does not match any other particulate or dissolved pool in the water column or sediments (Fig. 2c). Further, observed differences in δ^56^Fe between dissolved and particulate iron are consistent in magnitude and direction with those of other ferrotrophs in laboratory cultures (*19*). Although we cannot rule out the possibility of isotopic separation due to partial iron oxidation at the oxycline, the exceedingly low iron concentrations above the oxycline, the low water mixing rates between the metalimnion and hypolimnion and the high abundance of iron-reducing bacteria below and just above the oxycline imply that most or all accessible and re-oxidized iron should be reduced rapidly. This allows the hypolimnetic iron isotope signatures to be interpreted as the result of biological rather than chemical processes.

Our dissolved iron isotope composition profiles in the anoxic layer of L227 (and Lake 442) are distinct from those observed in Lake Pavin (*17*), which is a permanently stratified lake with an anoxic bottom water layer and hypothesized as an Archaean ocean analogue. The observed gradient of dissolved δ^56^Fe in Lake Pavin has been interpreted as resulting from partial oxidation of ferrous iron by O_2_ at the redox boundary (*17*). In addition, iron phosphates have been shown to be the predominant phase of the particulate matter in the anoxic monimolimnion (*20*). The total dissolved phosphorus (TDP) concentrations in both L227 and L442, up to 2 μM in the hypolimnion, are more than two orders of magnitude lower than that in Lake Pavin (up to 400 μM in the monimolimnion, *17*). Further, soluble reactive phosphorus concentrations are at or below detection limits (data not shown) implying that much of the TDP is in organic form. Thus, the formation of iron phosphates does not contribute to the observed water column δ^56^Fe in these lakes. In addition, photoferrotrophic bacteria have not yet been identified in association with minerals in the Lake Pavin water column (*20*). A lack of iron isotope compositions for the suspended particulates in Lake Pavin prevents further comparison between the two lake systems.

The similarity of δ^56^Fe in the hypolimnetic dissolved phase, epilimnetic POM and sediments (Fig. 2c), and the offset between hypolimnetic and epilimnetic POM (concomitant with δ^13^C and δ^15^N evidence), indicate that the δ^56^Fe signature in hypolimnetic POM results from microorganisms residing at that depth. Together, the microbial and geochemical data indicate that iron isotopes could be diagnostic of photoferrotropic activity in softwater lakes.

### Evidence from other Boreal Shield lakes

Photoferrotrophic activity is not confined to experimentally eutrophied L227. Similar geochemistry and microbiota were identified in unperturbed L442. Unlike L227, where the hypolimnion is entirely anoxic, L442 has both an anoxic zone and overlying oxic portion within the hypolimnion. Given low hypolimnetic mixing rates, this feature allows the stable isotopic signatures to be interpreted as a direct result of anoxia at similar temperature and organic matter supply. Natural abundance stable isotopes of C and N (Fig. 2) and water column chemistry (high Fe, DIC and NH_4_^+^, and low SO_4_^2-^; Fig. 1) in the anoxic hypolimnion of L442 are typical of anoxic hypolimnia in other ELA lakes. Patterns in particulate and dissolved phase δ^13^C, δ^15^N, and δ^56^Fe in L442 are all consistent with the anoxic hypolimnion of L227, and distinctly different from the oxic hypolimnion (Fig. 2). Molecular sequencing in the anoxic zone confirms high abundance of *Chlorobi* that are closely related to *C*. *ferrooxidans*, in addition to a similar microbial consortium of putative iron reducers, sulfur reducers and oxidizers, and methanotrophs (Fig. 3). Thus, even though boreal shield lakes are high in terrestrially derived organic carbon, unlike the Archaean Ocean, processes similar to the other reported Archaean analogues (e.g., *1, 3*) likely occur in Boreal Shield lakes with seasonally anoxic hypolimnia because the geology naturally leads to low sulfate and high iron conditions. Observations of both *Chlorobi* and methanotrophs have also been reported in anoxic hypolimnia in numerous Boreal Shield lakes in Finland (*21-23*). Further, in a recent study in Sweden, a shotgun metagenomics approach to the characterization of total DNA extraction was applied to a suite of Boreal lakes and indicated the presence of organisms having the ability to perform photoferrotrophy and anaerobic oxidation of methane (*24*). These new studies yield added support that these processes are widespread in Boreal Shield lakes.

## Discussion

Collectively, our results provide first evidence for potential photoferrotrophy in Boreal Shield lakes, showing how distinct isotopic and chemical indicators can be used to prospect for corresponding microbial consortia. Although conditions on present day Earth cannot totally mimic conditions in the Archaean Ocean, Boreal Shield lakes and ponds are globally abundant and have metabolically active processes relevant to Archaean Ocean microbial life. Moreover, new metabolic pathways associated with potential photoferrotrophy will also alter current paradigms of biogeochemistry in modern lakes and reservoirs.

Important and novel to the debate surrounding evolution of life in the Archaean Ocean is that the microbial consortia associated with potential photoferrotrophy in lakes are robust and establish rapidly. Both L227 and L442 have two mixing periods per year, with fall turnover being the most complete. In wind protected L227, the water column is well oxygenated every year in both spring and fall to at least 6 m where *Chlorobium* spp. are abundant (Fig. 3 and Supplementary Fig. S4). In L442, mixing is complete in both spring and fall (Supplementary Fig. S5). Isotopic and molecular analyses over several years show that these characteristic microbial consortia are re-established following oxygenation (Figs. 2 and 3). Although *Chlorobi* and methanotrophs have been found to re-establish in other lake hypolimnia following turnover (*22, 25*), our study is the first to show that the microbial consortia associated with photoferrotrophy are sufficiently resilient to rapidly recolonize after periods of oxygenation. The potential for anoxygenic photosynthesis in iron-rich systems that undergo periodic oxygenation may be relevant to understanding the formation of banded iron formations, especially immediately prior to the Great Oxygenation Event, where the relative roles of oxygenic and anoxygenic photosynthesis in iron oxidation and deposition remain controversial and oxygen-rich conditions were intermittent (*26*). Further, a strain of the filamentous cyanobacterium *Aphanizomenon schindlerii* was also observed at 6.5 m (*27*), leaving open the possibility that a hypothesized photoferrotrophic precursor to oxygenic photosynthesis may still be operating today under similar conditions to early anoxic oceans. That potential photoferrotrophs were detected in several small and nondescript Boreal Shield lakes in northern Canada, and likely also in Finland and Sweden, implies that these bacteria are more widespread than previously thought and are quite resilient. Boreal shield lakes provide new opportunities for whole ecosystem study of iron-cycling in ferruginous systems that may also provide constraints on early Earth processes.

Exploring the physicochemical conditions and isotopic fractionations associated with Boreal Shield lake anoxic zone microbial consortia will catalyze further scientific advances in both modern and ancient systems. We report first evidence of distinctive microbial fractionation of iron isotopes *in situ* in a setting with likely photoferrotrophic activity. Probing conditions in modern systems will facilitate new interpretations of the wide range of iron isotopic values observed throughout the Archaean (*28*). Similarly, our carbon isotope results have implications for both modern and ancient carbon cycling. Isotopic fractionation inherent in the reductive citric acid pathway (4 to 13 ‰; *29*), hypothesized as the photosynthetic pathway for GSB (*4, 6*) is too small to offset the much higher δ^13^C of the high flux of transitory organic matter and the associated heterotrophic activity in order to match the observed δ^13^C of the POM in the L227 hypolimnion given the measured δ^13^C of the DIC. Consumption of dissolved CH_4_ with very low δ^13^C, perhaps by the abundant methanotrophs in the anoxic zone, must contribute to L227 POM. Use of CH_4_ as a carbon substrate within such microbial consortia could alter interpretation of the δ^13^C signature for photosynthesis in ancient rocks and in modern settings, such as that published for Lake Kivu (*6*). In addition, biogenic CH_4_ is hypothesized to be an important contributor to the Archaean atmosphere (*30*). Thus, anaerobic consumption of biogenic CH_4_ also has implications for ancient carbon cycling. Finally, analyses of δ^15^N show evidence of isotopic fractionation when large amounts of NH_4_^+^ are present. Little is known about the expected ranges for δ^15^N in the Archaean Ocean. Although the observed shift in δ^15^N from negative values in the early Archean to more positive values in various geologic materials has been linked to the Great Oxidation Event, the interpretation remains controversial (*31*). Negative values in δ^15^N observed in early Archaean kerogens (*32*) could be attributed to metabolic fractionation of NH_4_^+^. However, the use of these kerogens as biosignatures has been challenged (*31*) and only laboratory experiments have been invoked to constrain the isotopic effects (*31*). Finally, large shifts in C, N, and Fe isotopic compositions have been recorded between ^~^2.8 and ^~^2.5 Ga ago and attributed to environmental or metabolic changes in the nascent period of early Earth oxidation (*29, 33*). Interpretation of the wide isotopic ranges preserved in different proxies in the geologic record would be aided by the modern study of these distinctive microbial consortia.

Our finding of active microbial iron oxidation in anoxic hypolimnia of Boreal Shield lakes also has novel implications for limnology, water management, and microbial ecology. Although bacterial iron cycling has been observed in the water columns of meromictic lakes (*34*), the overall importance of internal iron reduction-oxidation to metabolism within seasonally anoxic hypolimnetic water columns, including the presence of a putative iron oxidizer and high relative abundance and diversity of iron reducers, has not yet been recognized. Metabolism of gammaproteobacterial methanotrophs that are present in high abundance in the anoxic zone has been suggested to be coupled to oxygen produced by cyanobacteria at the same depth (*26, 35*), but it is also possible that this process may be coupled with reduction of newly oxidized iron by photoferrotrophs. Very recently, methanotrophs of the same taxonomic group as we found in L227 and L442 were found thriving in the anoxic zones of Lake La Cruz and meromictic Lake Zug, Switzerland and able to oxidize methane without oxygen in the absence of light (*14, 36*). Further, these methanotrophs were stimulated by the addition of oxidized Fe and Mn (*14, 36*). Iron reduction coupled to anaerobic methane oxidation has also been reported in an archaeal enrichment culture, showing the validity of this redox process in the natural environment (*37*). The importance of methanotrophy in anoxic hypolimnia with respect to either metabolism or CH_4_ emissions to the atmosphere is only starting to receive scrutiny (*25, 35, 38*), and not yet in the context of the metabolic pathways of the entire microbial consortium. Metagenomic data may shed light on the nature of anaerobic methane oxidation processes in Boreal Shield lakes and how these processes relate to those reported in other low-sulfate environments (*39*). Further, availability of reduced iron has recently been implicated as a factor controlling the dominance of cyanobacteria in both non-eutrophic systems and in the formation of hazardous algal blooms that have increased in frequency and severity in recent years (*27*); further understanding of iron redox cycling is urgently needed. Also unknown is the role of oxidized iron production in anoxic hypolimnia with respect to sequestration of phosphorous, the limiting nutrient in most Boreal Shield lakes. Lake 227 and similar boreal lakes have atypical low internal phosphorous release (*40*). Lastly, some iron cycling organisms are known to play a role in mercury (Hg) cycling and formation of methylated Hg (*41*), a toxic and bioaccumulating contaminant of fish in Boreal Shield lakes.

Using real systems is a powerful approach for promoting new discoveries. In contrast to simplified and hard to maintain laboratory cultures, whole thriving microbial communities can be studied under *in situ* environmental conditions, with additional layers of complexity that cannot be simulated in a laboratory. Boreal lakes and ponds number in the tens of millions (*42*) and, of these, some 15% could have a portion of the water column that is seasonally anoxic, opening new avenues of exploration. Water column chemistry and light penetration differ considerably among boreal lakes depending on geology and contribution of dissolved organic carbon from the catchment. Thus, broad gradients of physico-chemical conditions such as flux and quality of organic matter, sulfate and sulfide concentrations and light penetration can be exploited to better understand these unique microbial communities. Furthermore, small lakes, such as those at ELA, can be manipulated experimentally so that a specific range of conditions can be targeted or purposeful additions of carbon or iron isotope tracers can be used to probe isotopic fractionation *in situ*. Because hypolimnia become isolated during the stratified period and mixing is at rates of diffusion, accumulation or loss of constituents and mass balances can be used to infer metabolic pathways, products, and rates. Coupling these data to parallel metagenomic and metatranscriptomic analyses will facilitate reconstruction of genomes and active metabolic processes associated with photoferrotrophy. In addition, isotopic fractionations of δ^13^C, δ^15^N, and δ^56^Fe associated with anoxygenic photosynthesis and iron cycling that may be preserved in the global rock record can be studied *in situ* and during early stages of diagenesis in lake sediments deposited over the last 10,000 years since deglaciation. Boreal Shield lakes provide natural and accessible incubators for the study of processes relevant to the biogeochemistry and evolution of life in early Earth history.

## Methods

### Experimental design and site description

To assess the potential for the use of Boreal Shield lakes to yield insight into microbial Fe cycling processes, two exceptionally well characterized lakes (L227 and L442) were selected for detailed study over several years. These lakes are located at a long term scientific research site where comprehensive data on lakes, streams, climate, and hydrology have been collected for over 47 years, providing support for studies of shorter duration. Here, a suite of geochemical measurements, novel stable isotopic analysis, and nucleic sequencing provided complementary information on the occurrence of distinctive microbial consortia in these lakes.

The Experimental Lakes Area (ELA) is located in northwestern Ontario, Canada at 49°40’ N, 93°45’ W. Information on geology, vegetation, and climate is available (*43*). Lake 227 is a small headwater lake of 5 ha with a mean depth of 4.4 meters and maximum depth of 10 m. Lake 442 is a small lake of 16 ha with a mean depth of 9.6 meters and maximum depth of 17.8 m.

### Sample collection and analysis

Water and lake sediment samples from Lake 227 have been collected from the beginning of the nutrient addition experiment in 1969 and continue today. For this study, additional water, sediment trap, and sediment samples were collected over several years. Multiple water column profiles were collected in Lake 227 in 2010 and 2011 during the summer stratified period from May to October. Single profiles were collected around the time of the cyanobacterial bloom that occurs each year in June to July in 2012, 2013 and 2014. Exact sampling dates and parameters are given in Figures 1-2 and Supplementary Fig. S2.

Water samples were collected using a gear pump in a closed system from the desired depth. Samples for NO_3_^−^, NH_4_^+^, DOC, SO_4_^2−^, and total dissolved iron (TDFe) were filtered shortly after collection with 0.45 μm filters and analyzed using conventional methods (*44*). Sulfide was measured with an ion selective electrode (ISE) on unfiltered samples following stabilization in a sulfide anti-oxidant buffer (*32*). However, it is recognized that “free sulfide” is overestimated by an order of magnitude using the commonly used methylene blue method for dissolved samples due to the presence of “multiple reduced diffusible sulfur species” (*3*). The contribution of dissolved, colloidal and particulate sulfur species to the response by the ISE in our unfiltered samples is unknown but most likely causes a substantial overestimation of “free sulfide” species. Nitrate (NO_3_^-^) is very low in both the epilimnion and hypolimnion for these lakes with mean values of < 3.2 μM in the summer stratified period. The pH of these lakes varies between 6.2 to 6.8 throughout the water column Water column light levels are routinely measured at ELA in the ice-free seasons using a LICOR flat plate quantum sensor (LI-192) with a LI-250 light meter. The sensor measures PAR wavelengths (400-700 nm) with a limit of detection of 0.01 μE m^−2^ s^−1^. As a result of light scattering and absorption associated with high phytoplankton densities in L227, PAR is strongly attenuated over depth and only a very small proportion of surface irradiance reaches the hypolimnion. Throughout the May-October period, PAR values at 6 m depth in L227 were < 1.5 μE•m^−2^•sec^−1^. Detailed light profiles from L227 and L442 in June and September 2016 are presented in supplemental data (Supplementary Fig. S2).

Samples for concentrations and isotopic analysis of DIC and CH_4_ were collected directly without headspace into small glass serum bottles with stoppers and preserved with injections of concentrated HCl. Particulate organic matter (POM) for isotopic analysis was collected on Whatman QMA quartz filters with a nominal pore size of ~1 μm. Samples for microbial sequencing were collected by pumping water directly onto sterilized 0.22 μm Sterivex polyvinylidene fluoride filters (EMD Millipore). Filters were frozen until shipped back to the University of Waterloo and subsequently kept at −20°C or −80°C until processing.

Sediments were collected by freeze-coring (*46*) at 7 and 10 meters during the winter ice cover period at L227. The entire sediment profile was analyzed for δ^13^C and δ^15^N at 1 cm intervals. Selected samples were analyzed for δ^56^Fe. Additional sediment samples were also collected by subsampling an Eckman dredge at 1, 4, 8, and 10 meters in Lake 227 and at 1, 5 11, 13, 15, and 17 meters in Lake 442. The top 1 cm was collected from only those dredges where the surface was clearly visible and surface structures were undisturbed. Sediment traps constructed of acrylic tubes with a width to depth ratio of > 8 were suspended at 2.0, 5.5, and 8.5 meters depth in the water column at three locations in the central part of L227.

Samples for δ^13^C of DIC and CH_4_ were prepared by headspace equilibration after acidification. For analysis of δ^15^N of NH_4_^+^, samples were prepared using a modified diffusion technique (*47*) with a precision of ±0.3‰ in δ^15^N. Both δ^13^C of DIC and CH_4_ and δ^15^N of NH_4_^+^ were then analyzed by GCCF-IRMS using an Agilent 6890 GC coupled to an Isochrom isotope ratio mass spectrometer (IRMS: Micromass UK) with precision +/−0.3‰. δ^13^C, δ^15^N, and C/N of POM on filters, freeze-dried DOM, lake sediments, sediment trap samples were analyzed by EA-CF-IRMS using a Carlo Erba Elemental Analyzer (CHNS-O EA1108) coupled with a Delta Plus (Thermo) isotope ratio mass spectrometer with a precision of 0.2‰ in δ^13^C and 0.3‰ in δ^15^N.

Samples of POM, in water and sediment samples were analyzed for δ^56^Fe of Fe by MC-ICP-MS (Micromass Isoprobe) after purification using ion-exchange chromatography (*48*) and reported relative to the average of igneous rocks (δ^56^Fe = 0.0±0.05‰) with a precision of 0.03‰ (2σ). Sediment samples were digested with concentrated HF and HNO_3_ and then dried before loading onto the resin. The measured Fe isotope composition of the IRMM-019 Fe isotope standard was −0.08±0.05‰, which lies within error of the long-term value used in the lab of −0.09‰ relative to average igneous rocks (*48*).

### Microbial community analysis

Genomic DNA was extracted using the PowerWater Sterivex DNA Isolation Kit (MoBio) and quantified using agarose gel electrophoresis. The V3-V4 region of the bacterial 16S rRNA gene was amplified from each sample using triplicate PCR amplifications. Each reaction contained ≤10 ng of sample DNA and used reagent, volumes, and thermocycler conditions described previously (*49*). Modified forward and reverse PCR primers 341f-808r were used for Illumina sequencing in a previously described configuration (*50*). After combining triplicate reaction products to reduce bias, products for each sample were pooled at a normalized concentration, and the pooled library was gel purified using the Wizard SV Gel and PCR Clean-Up System (Promega), and spiked with 8.5-10% PhiX prior to sequencing. Paired-end (2x250 base) high-throughput DNA sequencing was carried out using the MiSeq platform (Illumina), achieving a cluster density of 452-507 K mm^−2^ with 92.9-98.0% of clusters passing filter. An average of ~ 600,000 raw reads per sample were generated for downstream analysis. Raw demultiplexed sequencing reads were processed using the AXIOME2 software tool, version 1.5 (*51*). Using AXIOME2, paired sequences were assembled using PANDAseq version 2.8 (*52*), were chimera checked and clustered at 97% using USEARCH version 7.0.1090 (*53*), and were rarefied using QIIME version 1.9.0 (*54*).

To predict the major functional roles of the lake bacterial communities, the top ten most abundant bacterial OTUs within each water column sample, based on rarefied data, were evaluated for their potential involvement in Fe or S oxidation or reduction, and methanotrophy. Representative sequences for each abundant OTU were assigned taxonomic ranks using the RDP Naïve Bayesian rRNA Classifier, version 2.10, with a confidence threshold of 50%, based on the RDP 16S rRNA training set 14 (*55, 56*). If the classifier could not assign a rank at the genus level, OTU representative sequences were queried against the NCBI non-redundant nucleotide database using BLASTN (*57*). Cultured strains identified through the query with ≥97% sequence identity were used to assign, where possible, a single genus to the OTU. Subsequently, genera matched to each abundant OTU were checked against the literature for whether they contained strains with documented involvement in targeted metabolic activities. Any OTU classified within a genus that was found to contain a strain involved in one of these metabolic activities was inferred to have the same potential metabolic activity. Potential metabolic functions of OTUs that could only be classified to the family level were inferred similarly. Any remaining OTUs were not assigned a metabolic role.

High abundance OTUs closely related to the photoferrotroph *C. ferrooxidans* were also examined phylogenetically to evaluate their placement within the green sulfur bacteria. Reference 16S rRNA gene sequences of cultured strains from the family *Chlorobiaceae*, along with an appropriate outgroup sequence, were obtained from the Silva SSU-RefNR sequence database, release 122 (*58*). Obtained sequences were aligned to representative sequences of each abundant *Chlorobium* OTU from L227 and L442 using SINA version 1.2.11 (*59*), and the alignment was then truncated to the V3-V4 region of the 16S rRNA gene. Using this alignment, a phylogeny was built using RAxML version 8.1.17 (*60*), using 100 maximum likelihood searches and the GTRCAT sequence evolution model. Node support values were calculated using the Shimodaira-Hasegawa test. The resulting phylogeny was visualized using Dendroscope version 3.2.10 (*61*).

Raw amplicon sequencing data is available in the NCBI sequence read archive under Bioproject [accession numbers to be inserted once sequence submission is complete]. Representative sequences for each OTU, determined using AXIOME2, were deposited in the NCBI under accession numbers [accession numbers to be inserted once sequence submission is complete]. Counts of OTUs in each lake water sample are available in Supplementary Data File S1 online as an OTU table.

## Acknowledgements

We thank researchers at the Experimental Lakes Area who amassed an unparalleled dataset over the past 46 years, providing crucial data in support of shorter term studies. We thank D.W. Schindler, R.E. Hecky, W.D. Taylor, and C. Welte for critical reading of the manuscript, S. McCabe and staff at the Experimental Lakes Area for technical help with chemical analyses, K. Liu for δ^56^Fe analysis, C. Johnson and B. Beard for providing the facility for δ^56^Fe analysis, and K. Engel for assistance with sequencing. All authors were funded by the National Sciences and Engineering Research Council of Canada (NSERC) and the Water Institute at the University of Waterloo, Canada.

## Author contributions

SLS, JJV, LM, MP were involved in study design of the larger project that supported data collected for this paper. SLS, JJV, RJE collected samples and analyzed δ^13^C and δ^15^N. LW analyzed samples for δ^56^Fe. JMT and JDN performed molecular analysis. SLS wrote the paper with JDN, JMT, LW and comments from all other authors.

## Author Information

Correspondence should be addressed to SLS or JDN.

## SUPPORTING INFORMATION

**Supplementary Figure 1.**
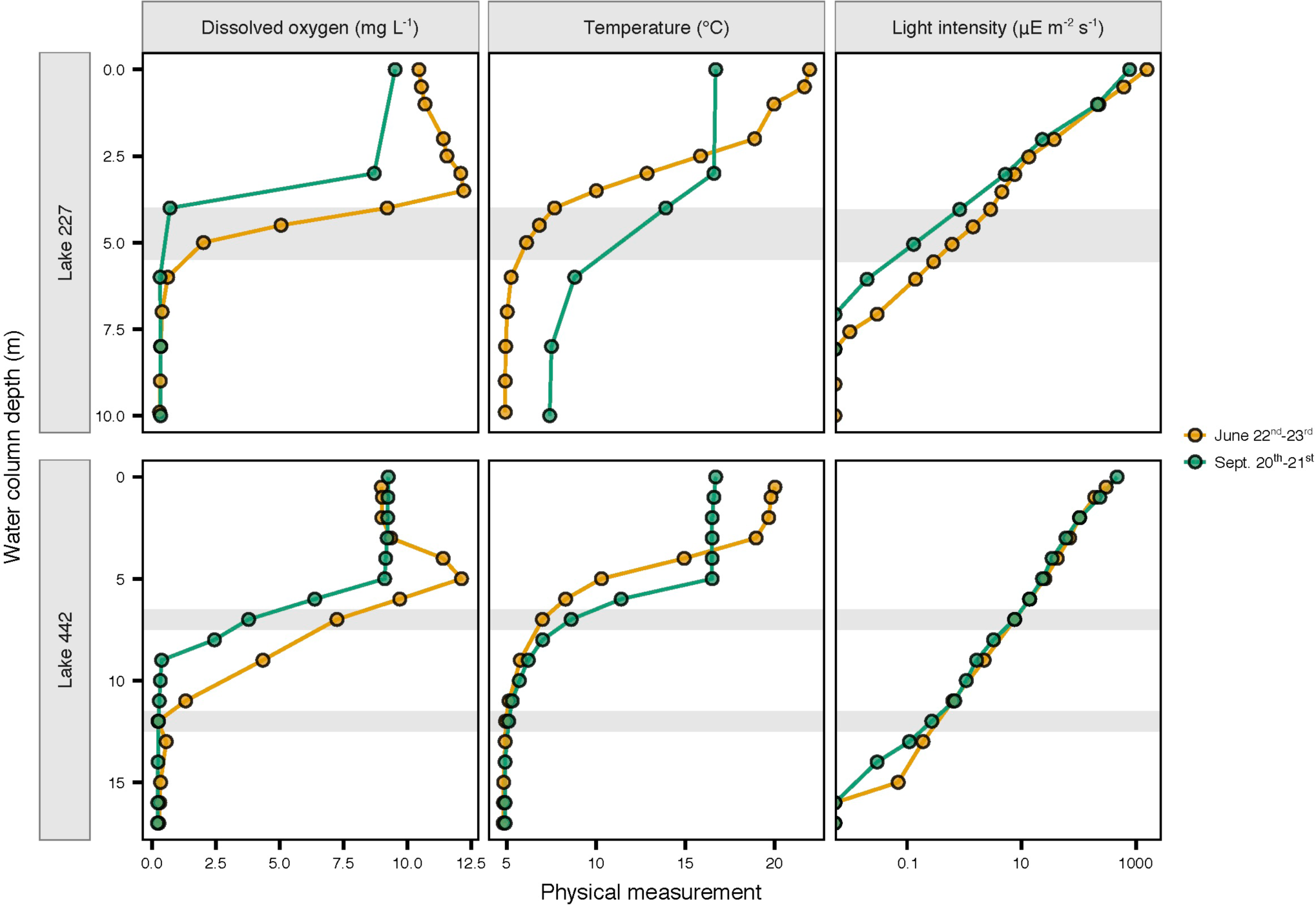
Dissolved oxygen, temperature and light profiles of the L227 and L442 water columns in 2016. Grey bars represent approximate transition zones between lake layers and are identical to those shown in Figs. 1-2 (see figure captions). Exact layer boundaries at the sampling times may be estimated using the temperature and dissolved oxygen profiles shown. In both lakes, low but detectable light levels reached the top of the anoxic zone in both June and September. Light measurements below the detection limit (0.01 μE m^−2^ s^−1^) are plotted along the left edge of the y-axis for reference.

**Supplementary Figure 2.**
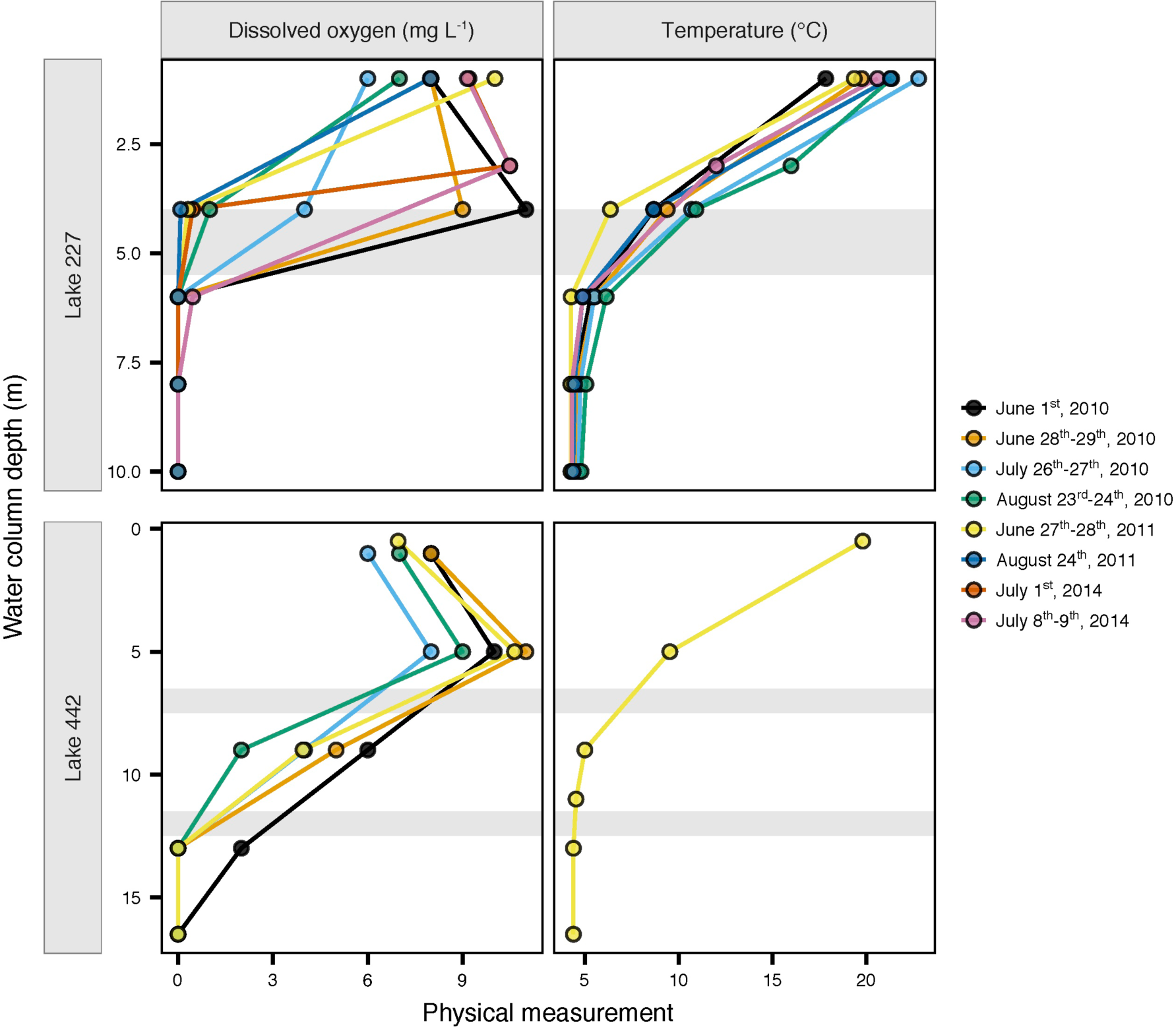
Physical profiles of the L227 and L442 water columns during isotopic and molecular sampling. Grey bars represent approximate transition zones between lake layers and are identical to those shown in Figs. 1-2 (see figure captions). For comparison, actual lake zone boundaries at each time of measurement may be determined visually from temperature and dissolved oxygen data. Seasonal variations can be noted in the temperature and oxygen status of the water columns at depths near the approximate transition zones, for example for L227 at 4 m depth.

**Supplementary Figure 3.**
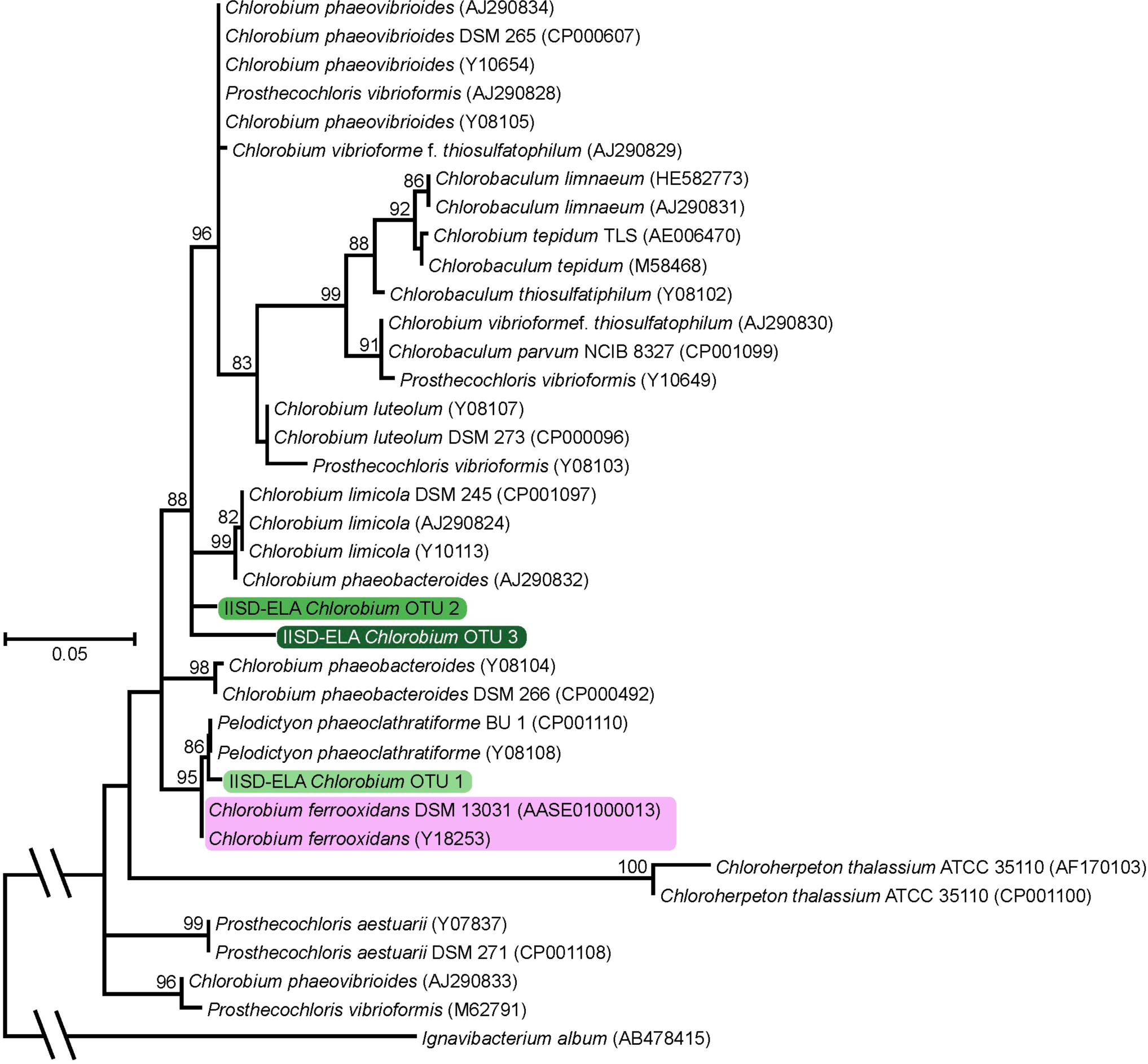
Phylogenetic placement of potential photoferrotrophs within the family ***Chlorobiaceae***. Reference *Chlorobiaceae* sequences represent cultured strains. Node support values, calculated using the Shimodaira-Hasegawa test, are shown where 80% or higher. Sequences highlighted in pink correspond to known photoferrotrophs, and those highlighted in green represent OTUs identified at high abundance in the water columns of L227 and L442 (Fig. 3). Importantly, IISD-ELA *Chlorobium* OTU 1 was identified at high abundance in the water columns of both lakes.

**Supplementary Figure 4.**
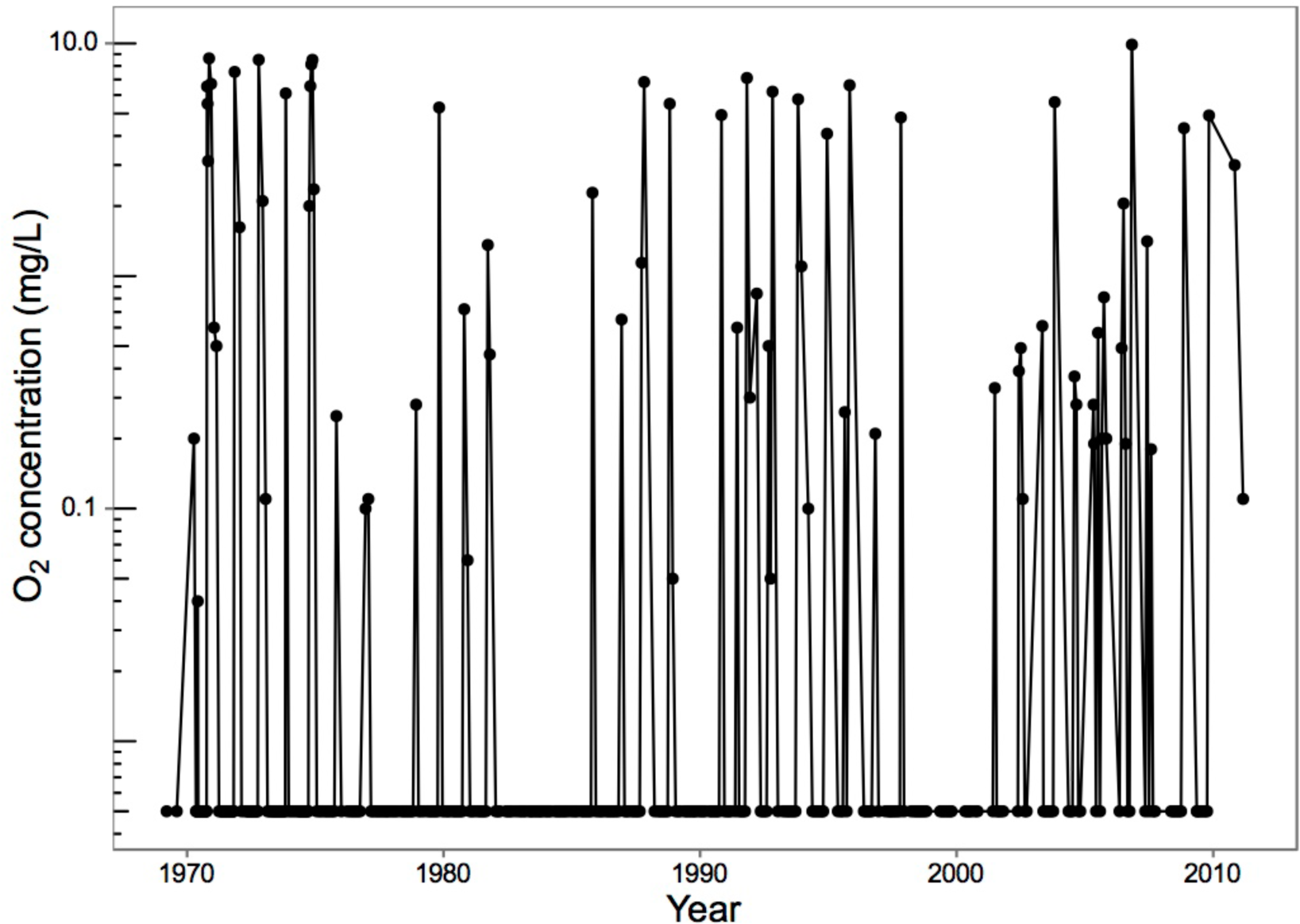
Dissolved oxygen at 6 m depth in the L227 water column from 1969 to 2011. The detection limit for O_2_ is 0.005 mg O_2_ L^−1^. *Chlorobium* sequences were detected at high abundance in both 2013 and 2014 at this depth (Fig. 3, please also see legend). Dissolved oxygen samples were collected typically at least every two weeks in summer but were collected at most twice during the winter. Following this sampling schedule, the full extent of the typical spring and fall re-oxygenation events (overturns) in L227 may not have been measured in some years. Dissolved oxygen is typically, but not always, measured after fall overturn. Spring overturn measurements can be missed following ice-off due to logistical reasons and especially in years when temperatures warm rapidly after ice-off. Thus, the oxygen record at 6 m reflects the minimum number of re-oxygenation events at this depth.

**Supplementary Figure 5.**
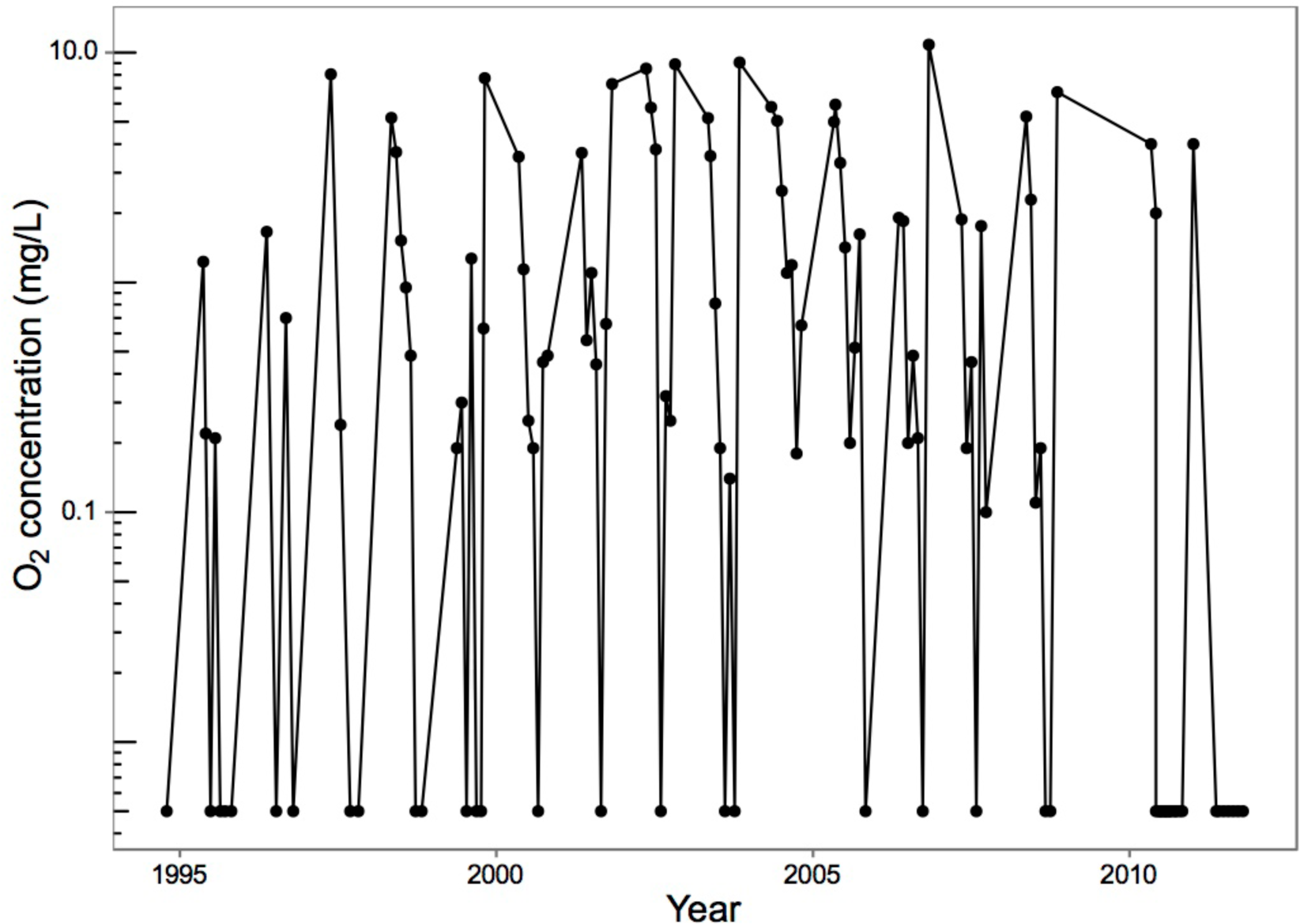
Dissolved oxygen at 13 m depth in the L442 water column from 1994 to 2012. The detection limit for O_2_ is 0.005 mg O_2_ L^−1^. *Chlorobium* sequences were detected at high abundance at this depth in both 2011 and 2014 (Fig. 3). Dissolved oxygen samples were collected typically once per month in summer but were collected at most once during the winter. Following this sampling schedule, the full extent of the typical spring and fall re-oxygenation events (overturns) in L442 may not have been measured in some years. Dissolved oxygen is typically, but not always, measured after fall overturn. Spring overturn measurements can be missed following ice-off due to logistic reasons and especially in years when temperatures warm rapidly after ice-off. Thus, the oxygen record at 13 m reflects the minimum number of re-oxygenation events at this depth.

**Supplementary Data File S1. Rarefied hit counts of operational taxonomic units (OTU) within water column samples of L227 and L442**. The OTU table (CSV format) was prepared using the software tool AXIOME2 (see Materials and Methods). Representative sequences of each OTU are included. Classification of each OTU is shown up to the taxonomic rank where the RDP classifier’s confidence value fell below 50%.

